# Meditation as a Bioactive Intervention: Molecular and Neurophysiological Mechanisms Revealed by Connectivity Mapping

**DOI:** 10.64898/2025.12.27.696659

**Authors:** Aditi Joshi, Deep Patel, Vignesh Muralidharan, Mitali Mukerji

## Abstract

Meditation practices are often used as non-pharmacological adjunct therapy for managing stress and well-being benefits. We propose that beneath their non-pharmacological facade, meditation practices might operate via drug target modulation having profound neurophysiological effects. Firstly, we leverage the Connectivity Map (CMap) to investigate (a) the overlap between meditation-induced molecular signatures and established drug responses, and (b) the pathways and mechanisms contributing to meditation’ potential therapeutic effects. This was studied in a comprehensive temporal RNAseq dataset comprising premeditation, meditation, and follow-up stages from a clinical trial involving 106 participants practising inner engineering meditation. Meditation signatures overlapped with over 438 drugs, but predominantly with drugs targeting the neuroactive ligand receptor pathways, capable of modulating the brain’s excitation-inhibition (E/I) balance. Next, to bridge the meditation-induced molecular changes to measurable neurophysiological effects, we performed a comprehensive meta-analysis of drugs affecting E/I balance through the lens of non-invasive brain stimulation, i.e., transcranial magnetic stimulation (TMS-EMG and TMS-EEG). Using multiple correspondence analysis, we then clustered the CMap informed neuroactive drugs and their E/I effects on the same latent space. We found that meditation’s effects mimic drugs targeting the GABAergic, Glutametergic and other neuromodulatory system pathways. Our findings lead to a working model to objectively test neurophysiological changes resulting from meditation that can lead to evidence-based clinical applications. Overall, these findings support the view that meditation acts as a biologically active intervention capable of modulating molecular pathways and cortical excitability, rather than functioning solely as a psychological or contemplative practice.

## Introduction

Meditation methods, which have their roots in Indian medicine and Buddhism, are used as non-pharmacological adjunct therapies for managing stress, sleeplessness, anxiety, pain, depression, and cognitive improvement (1). Meditation promotes rest, relaxation, and digestion by stimulating the parasympathetic nerve system, which counteracts the sympathetic “fight-or-flight” response(2). However, the physiological effects differ between individuals and are altered by procedures and durations, possibly resulting in a range of outcomes(3,4). Meditation might aggravate symptoms in people who already have mental health issues (ref - cheetah house related publication).

The biological effects of meditation have been explored from different perspectives. For example, findings from non-invasive neurophysiology, particularly transcranial magnetic stimulation (TMS) and a broad range of neuroimaging studies have revealed that meditation and other mind–body practices are known to influence brain function (5,6). Neuroimaging studies further indicate that meditation alters functional connectivity across major intrinsic brain networks, reflecting large-scale reorganization of circuits involved in attention, interoception, and emotion regulation(7). More recently, an intervention that combined meditative practice, cognitive reframing, and open-label placebo treatment demonstrated tightly coupled neural and molecular responses including short-term remapping of functional networks, enhanced markers of neuroplasticity, and coordinated changes in plasma transcriptomic and metabolic profiles. Such evidence supports the view that meditation influences not only cortical dynamics but also downstream biological pathways(8).

A recent large scale transcriptome study on advanced inner engineering meditation reported modulations of networks that are perturbed in severe COVID-19 infection and multiple sclerosis. This suggests meditation may be useful in conditions of compromised immune systems and increased inflammation(9). We first hypothesized that the molecular targets underlying meditation’s response might overlap with those modulated by pharmacological interventions. We used connectivity map (CMap) to query intersections of meditation response with over 3 million gene expressions of 5,000 small compounds and 3,000 genetic perturbations across 9 cell lines(10–12). Our study revealed convergence with over 438 pharmaceutical drugs, which were strongly connected to neuroactive ligand receptor signalling pathways. We then hypothesized that the influence of meditation, if similar to neuroactive drug action, will lead to similar changes in the cortical excitation–inhibition (E/I) balance and can be read out through non-invasive brain stimulation methods. One way to investigate the changes in E/I dynamics is the use of TMS along with muscle (TMS-EMG) or electroencephalography (TMS-EEG) readouts. The influence of specific drugs on the cortical excitability has been studied, with specific emphasis on those involving the GABAergic and glutamatergic neurotransmitter systems. Pharmaco-TMS work has demonstrated that indices such as motor threshold, cortical silent period, short-interval intracortical inhibition (SICI), and intracortical facilitation (ICF) are sensitive to changes in GABAergic and glutamatergic signaling. Pharmacological agents that act on these neurotransmitter systems consistently produce measurable shifts in cortical excitability in healthy individuals(13,14).

Motivated by this idea, we compiled and analyzed literature on pharmacological studies in which the effects of different drugs on cortical excitation–inhibition (E/I) balance were directly tested using TMS-EMG/EEG measures. Comparing the meditation-related transcriptional signatures with drugs and drug classes identified through CMap, alongside drugs known from the neurophysiology literature to alter cortical excitability we found converging evidence that meditation may act on neurobiological pathways similar to those influenced by pharmacological manipulation of the E/I system. The E/I balance reflects the interplay between glutamatergic excitation and GABAergic inhibition, and are closely linked to changes in motor output, sensory integration, and attentional control. This supports the idea that meditation, although non-pharmacological, can produce quantifiable and mechanistically meaningful changes in brain function. This systematic mapping enabled us to identify drug classes with well-characterized effects on motor-system physiology, providing a framework for interpreting how meditation may modulate E/I balance in the brain. The combined insights suggest that meditation practices may benefit from structured, individualized approaches to maximize effectiveness and safety.

## Materials and Methods

### 1. Study Cohort and Data Collection

The study was carried out on 389 transcriptomes (GSE174083), from 106 individuals of a meditation retreat (ClinicalTrials.gov identifier: NCT04366544). The RNAseq data were before the retreat (T1),after an 8-week preparatory phase (T2), 8 days after the meditation retreat (T3), and 3 months after the retreat (T4). Trimmomatic (version 0.39) with default parameters was used on raw reads for adapter removal and trimming(15) followed by STAR for mapping the RNA-Seq data, and DESeq2 (1.40.2) for identification of differentially expressed genes (log2fold changes 1 and p-value 0.05). To explore inter sample variability, the raw count of samples from T2 and T3 timepoints were TMM normalised prior to DESeq2 (1.40.2) and clusters were identified through hierarchical clustering (Scipy version 1.11.3) and Principal Component Analysis (Sklearn version 1.3.2).

### 2. Connectivity Map Analysis to identify interventionable points

The ranked lists of differentially expressed up and down regulated genes in pairwise comparisons of T2vsT3, T2vsT4, and T1vsT2 were used to query the library of gene expression profiles from connectivity map (CMap) data base. Signatures with a connectivity score of $\geq$ 98 were considered. Gene Ontology analysis was performed on the targets to uncover the enriched pathways and targets influenced by meditation-related gene signatures. Specifically, for inferring pathways and targets similar to those affected by drugs, the enrichment was performed using gene sets exhibiting positive connectivity with drugs and knockdown genes. Conversely, pathways aligning with drugs showing negative connectivity or corresponding to overexpressed genes were deduced as pathways exhibiting opposite effects.

### 3. Meta Analysis

The dominance of neuroactive ligand receptor interaction pathways temporally prompted further investigation into the effects of these drugs. A systematic literature search was conducted using PubMed and Google Scholar with the keywords *“Drugs AND TMS AND excitability.”* From PubMed, 125 articles were identified, of which 42 met the inclusion criteria. From Google Scholar, the top 100 search results were screened, yielding 25 relevant studies. The full PRISMA flowchart is provided in Supplementary Fig. S2.

Articles were considered relevant if they reported the effects of drugs on TMS-EMG or TMS-EEG measures, or if they discussed physiological changes induced by pharmacological agents. Review articles describing drug-related modulation of cortical excitability or neurophysiological processes were also included. Studies were excluded if they: (i) only used TMS as a therapeutic tool for psychiatric or neurological disorders without assessing drug effects, (ii) described excitability measures unrelated to pharmacological modulation, or (iii) lacked direct discussion of drug-induced physiological alterations. The detailed inclusion and exclusion process is summarized in Figure S2.

The following metrics were considered as the outcome measures for the drug effects on the brain as probed using TMS. For cortical excitability, measured using TMS-EMG, we looked at the motor system physiology metrics, i.e., the resting motor threshold (MT), motor-evoked potential (MEP) amplitude, intracortical inhibition measures including short-intracortical inhibition (SICI) and long-intracortical inhibition (LICI), intracortical facilitation (ICF), short afferent inhibition (SAI) and the cortical silent period (CSP). We also looked at cortical plasticity effects measured using paired associative stimulation (PAS). In case of the TMS-EEG metrics, we considered the typical TMS-evoked potential (TEP) positive peaks observed from P15/P15-25 complex, P60, P75, P100, P150 and P180 and negative peaks N45, N75 and N100. We also considered commonly seen TMS-evoked spectral perturbations (TERSPs) including early (30-200ms) alpha and beta synchronization, and late (200-400ms) alpha and beta desynchronization.

### 4. Integration of TMS Metrics with Connectivity Map Data

The meta-analysis provided a detailed table (Supplementary Table S5), summarizing TMS metrics, drug responses, and their respective classes. In parallel, we have pathway perturbations associated with meditation across time points through ontology-based analysis of Connectivity Map (CMap) data. For each pathway, the corresponding drugs and their classes were extracted from the CMap output. A comparative analysis of these two datasets was then performed to identify overlapping patterns that could indicate meditation-induced modulation of excitation–inhibition (E/I) balance via drug-related mechanisms and classes.

### 1. Data Preparation

A consolidated dataset was generated combining all drugs, their classes, TMS metric responses, and temporal CMap data (Supplementary Tables S6–S8). The two datasets were merged based on shared drug classes rather than individual drugs. The merged data was sparse, as not all drugs had corresponding TMS parameters reported, nor all classes from the CMap dataset had TMS parameters. In total, the dataset included 70 drugs, 28 unique drug classes, and 21 TMS parameters. For the CMap data, both positive and negative connectivity scores were included across three time points (T1 vs T2, T2 vs T3, and T2 vs T4). The merged dataset was coded such that drug classes from the TMS dataset appearing in the CMAP data were assigned a value of “1” present and “NA” if absent.

### 2. Multiple Correspondence Analysis (MCA)

To reduce dimensionality and visualize categorical associations, Multiple Correspondence Analysis (MCA) was applied. MCA is suitable for datasets comprising qualitative variables such as increase/decrease, present/absent, or categorical drug responses. Prior to MCA, drug classes were hierarchically consolidated into broader super classes (e.g., “dopamine receptor agonist,” “dopamine receptor antagonist,” and “dopamine modulator” were grouped as “Dopaminergic”). This reduction was performed for most drug classes (Supplementary Table S9), with remaining categories grouped under “Others.” MCA was implemented in R, where categorical responses were converted to factors, super classes were assigned, and analysis was performed using the MCA() function. Separate MCA analyses were conducted for each of the three time points.

The drug super classes were assigned based on the broad pharmacological systems including neurotransmitter-related classes (GABAergic, cholinergic, dopaminergic), ion-channel modulators, and metabolic or hormonal modulators. These groups capture the primary physiological system each drug acts upon, not the exact molecular mechanism (agonist, antagonist, blocker, etc.)(Supplementary table).

### 3. Cluster Analysis

Following MCA, K-means (K=3) clustering was applied to the MCA data to identify distinct groupings of drug classes based on their TMS response profiles and CMap connectivity patterns. Since MCA transforms categorical data into continuous coordinates, it enables effective distance-based clustering using K-means. Clusters with at least 3 drugs were calculated. The cluster centroids were identified, and the corresponding superclasses were mapped to CMap data for both positive and negative connectivity.

### 4. Cluster Stability Assessment

To evaluate the robustness of the identified clusters, a non-parametric bootstrap analysis was performed. The dataset was repeatedly resampled (n=300) with replacement, followed by MCA and K-means clustering for each bootstrap iteration. The similarity between cluster assignments from the original data and bootstrap replicates were quality assessed using the Adjusted Rand Index (ARI). ARI values close to 1 indicate high cluster reproducibility, while values near 0 denote random correspondence. The other statistical parameters to assess the cohesion and non- randomness of clustering, the Hopkins score, mean silhouette score and Fisher’s exact test were performed.

### 5. Direct overlap

We also identified direct overlaps between drugs showing TMS responses and those exhibiting significant connectivity with meditation-associated pathways across different time points. These overlaps were visualized using a Sankey plot to illustrate how specific drug classes intersect with distinct meditation time points. This visualization enabled identification of the dominant drug classes contributing to each time point and provided insights into their corresponding TMS response patterns, thereby suggesting potential temporal trajectories of cortical modulation during meditation.

## Results

We leveraged the demonstrated capabilities of the connectivity map, to discern the potential pharmacological connections during meditation (11). This was tested using RNAseq-based transcriptomes from a clinical study involving 106 participants in an advanced inner engineering meditation retreat program(9). The time course data, spanning from the initial phase (T1) through the 8-week preparatory phase (T2), the 8-day inner engineering retreat (T3), and a 3-month follow-up (T4), enabled us to explore specific effects. We then conducted a meta-analysis of studies investigating the effects of pharmacological agents on the E/I balance, as measured via the effect of non-invasive brain stimulation via TMS on motor system excitability. Following the meta-analysis, we aimed to integrate and map two axes of modulation relevant to meditation i.e. pharmacological and neurophysiological, to better understand meditation-induced alterations.

### Connectivity Analysis of Meditation Transcriptome reveals Potential Drug Targets

CMap revealed a significant intersection with over 438 drug signatures across the three time spans (Figure 1a, Supplementary table S1). There was a significant overlap in profiles of drug signatures in the preparatory and the meditation phase (T1vsT2, T2vsT3) which was very distinct from the follow-up phase (T2vsT4). The drug classes identified were diverse, encompassing cyclooxygenase inhibitors, serotonin receptor agonists/antagonists, glucocorticoid receptor agonists, dopamine receptor antagonists, HDAC inhibitors to name a prominent few (Supplementary table S2). Gene ontology (GO) analysis of the drug targets (a) drugs with positive scores and knockdown genes and (b) negative scores and over-expressed genes highlighted the enriched pathways through which meditation could exhibit its effects (Figure 1b, Supplementary table S3). Noteworthy, the neuroactive ligand receptor interaction pathway (hsa04080) consistently featured within the positive connectivity set across all three phases of meditation. The targets were quite distinct in the follow-up phase and corroborates with the patterns from drug signatures(Supplementary table S2). We followed this analysis with specific emphasis on the neuroactive drugs and their effects on the altering E/I balance in the brain.

**Figure 1:**
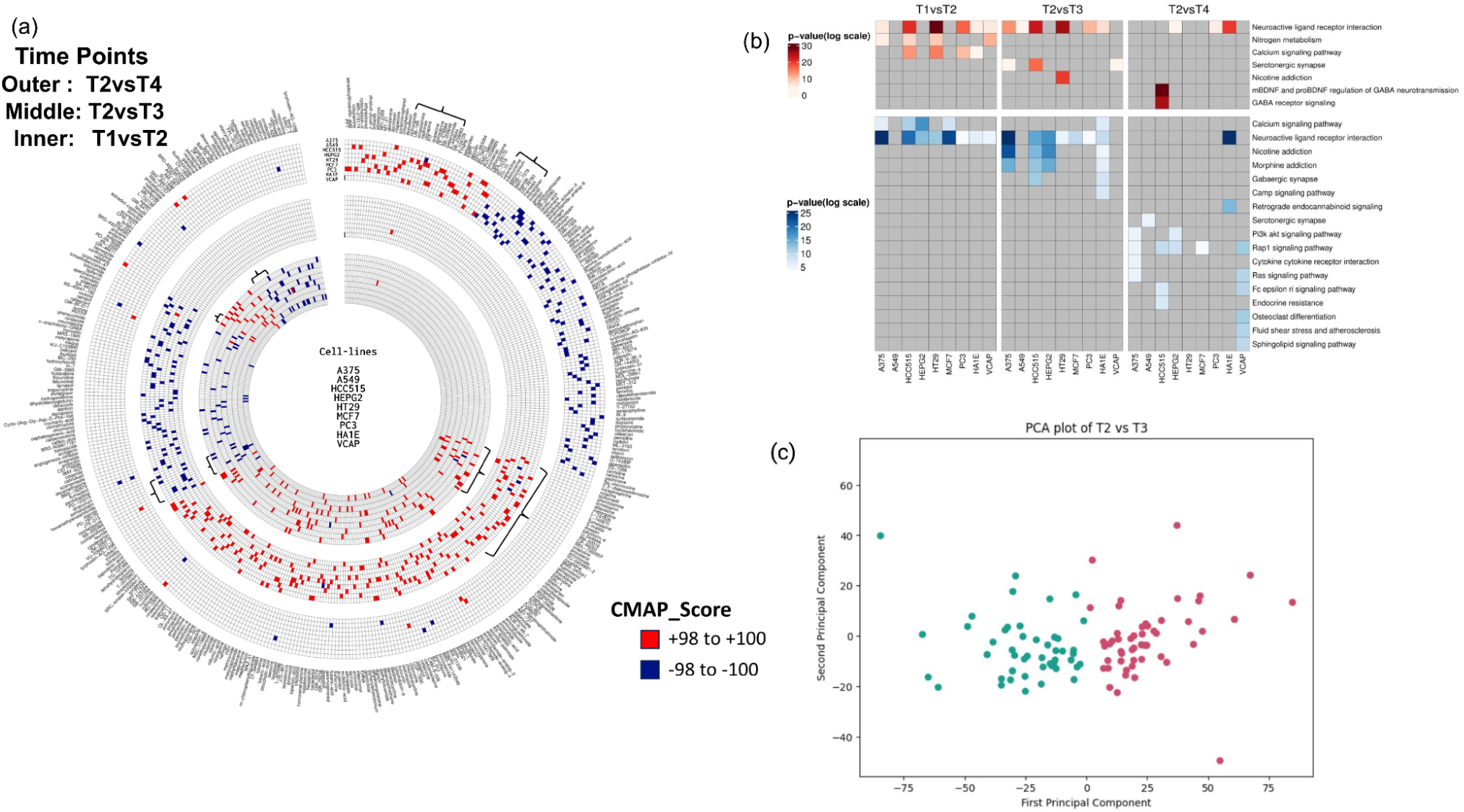
(a) Circos Plot of Connectivity Map Matching Signatures.The Circos plot depicts the Connectivity map analysis results for matching signatures of drugs across three distinct time points within subjects who participated in the advanced inner engineering meditation retreat. The time points under consideration are: before the retreat (T1), after an 8-week preparatory phase (T2), 8 days after the Samyama meditation phase (T3), and 3 months after the retreat (T4). Each concentric circle represents a specific comparison: the innermost circle corresponds to the connectivity of drugs with differential expression signatures from T1 vs T2, the middle circle represents T2 vs T3, and the outermost circle corresponds to T2 vs T4. In the visualization, drugs exhibiting connectivity with positive scores equal to or exceeding 98 are color-coded in red, while those with negative scores are depicted in blue. Remarkably, there is a substantial overlap of drug signatures between the preparatory and meditation phases. Notably, the number of drugs displaying connectivity in T2 vs T3 (243) surpasses that of the preparatory phase (210). Interestingly, after 3 months, a majority of these initial signatures dissipate, giving way to the emergence of new drug signatures. The drugs targeting the significantly enriched neuroactive ligand receptor signaling pathway are denoted by the symbol (Supplementary table S2).(b) Pathway enrichment analysis of proteins targeted by drugs across all three phases. The upper panel, highlighted in blue, showcases significant pathway enrichments based on Log p values derived from drugs with positive scores. The lower panel depicts enrichments from drugs with negative scores.(c) The PCA plot of sample wise expression difference in T2 vs T3 reveals two distinct clusters of meditation response within the study cohort (Supplementary Figure S1). Image created with BioRender.com.

### Inter-individual variability in meditation response

The inter-individual variability in meditation response was also an interesting observation. Intersection with such a large number of drugs could be a cumulative effect of differential response of individual samples. To test this, we carried out a sample-wise TMM normalization followed by differential expression analysis between T2 and T3 (P <= 0.05), PCA and hierarchical clustering (Figure 1c). Two distinct clusters of samples separated out in PCA that had distinct non-overlapping connectivity with drugs (Supplementary Figure S1). This shows that rather than eliciting a homogeneous biological effect meditation gives rise to distinct individual level molecular response patterns, which collectively manifest as broad drug associations at the cohort level.

### Drugs influencing E/I balance in the motor system can provide crucial insights in meditation practices

The observed bidirectional relationship between meditation and neuropsychiatric-related drugs in the Connectivity Map highlights meditation’s potential to modulate neural excitability and signaling. Motivated by this observation, we sought to explore the potential effects of these drugs on brain activity. If the effects of meditation are reflected as changes in the neuroactive receptor pathway systems, then understanding the changes in the cortical E/I balance in pre and post meditation can act as markers to provide evidence for meditation as a bioactive intervention. To evaluate this possibility, we decided to map to a common space, the CMap profiles and the drug-induced neurophysiological changes.

To do this we first did a comprehensive meta analysis of drugs affecting cortical E/I balance as read out using TMS metrics (Supplementary Figure S2) (for details see Methods section: Meta-Analysis). This revealed distinct relationships among several meaningful TMS–EMG/EEG metrics detailed in Supplementary Table S5 for all the drugs and their TMS interactions. To highlight a few interesting relationships, there were drug classes that predominantly increased GABAergic inhibition, which was reflected by reduced motor-evoked potential (MEP) amplitudes, suppression of TMS-evoked potentials (TEPs) and enhancement of intracortical inhibition (short and long intracortical inhibition, SICI and LICI). There were drugs affecting glutamatergic transmission including NMDA receptor antagonist which increased SICI and decreased intracortical facilitation. Drugs acting through the monoamine neurotransmitter systems, including serotonergic, dopaminergic and norepinephrinergic systems, led to a heterogeneous pattern of changes. Dopamine agonists influenced changes in cortical plasticity, inducing long-term potentiation, as measured using paired-associative stimulation (PAS). Norepinephrine reuptake inhibitors in general increased MEP amplitude, decreased SICI and increased intracortical facilitation (ICF). From this detailed relationship, we wanted to understand how meditation would bring about neurophysiological changes by mapping both the CMap derived drug information and the drug-related TMS effects.

To address this, we then integrated this data with the Connectivity Map (CMap) perturbational profiles to explore how meditation influence E/I balance through pharmacological analogies. The Multiple Correspondence Analysis (MCA) followed by clustering based in MCA space, identified temporal patterns that reflect how neural excitability shifts (Figure 2). In the T1vsT2 (preparatory) phase (Figure 2a), clusters showed representation of GABAergic and dopaminergic drug classes, consistent with early dominance by inhibitory and reward-associated mechanisms. Additional drug classes showing positive or negative connectivity included endocannabinoid, glutamatergic, and serotonergic groups, aligning well with initial neurochemical adaptations during initial meditation practice and dietary change. During the T2vsT3 (Inner Engineering) phase (Figure 2b), the co-clustering of GABAergic drugs with calcium channel modulators, with the cluster centroid aligning closely with calcium-associated profiles. This convergence suggests a shared multivariate signature consistent with concepts central to maintaining cortical E/I balance. Prominent drug classes included glutamatergic, dopaminergic, adrenergic, and serotonergic, indicating active engagement of both inhibitory and excitatory neurotransmitter systems, suggesting behavioural, emotional control and modulation towards plasticity.

**Figure 2:**
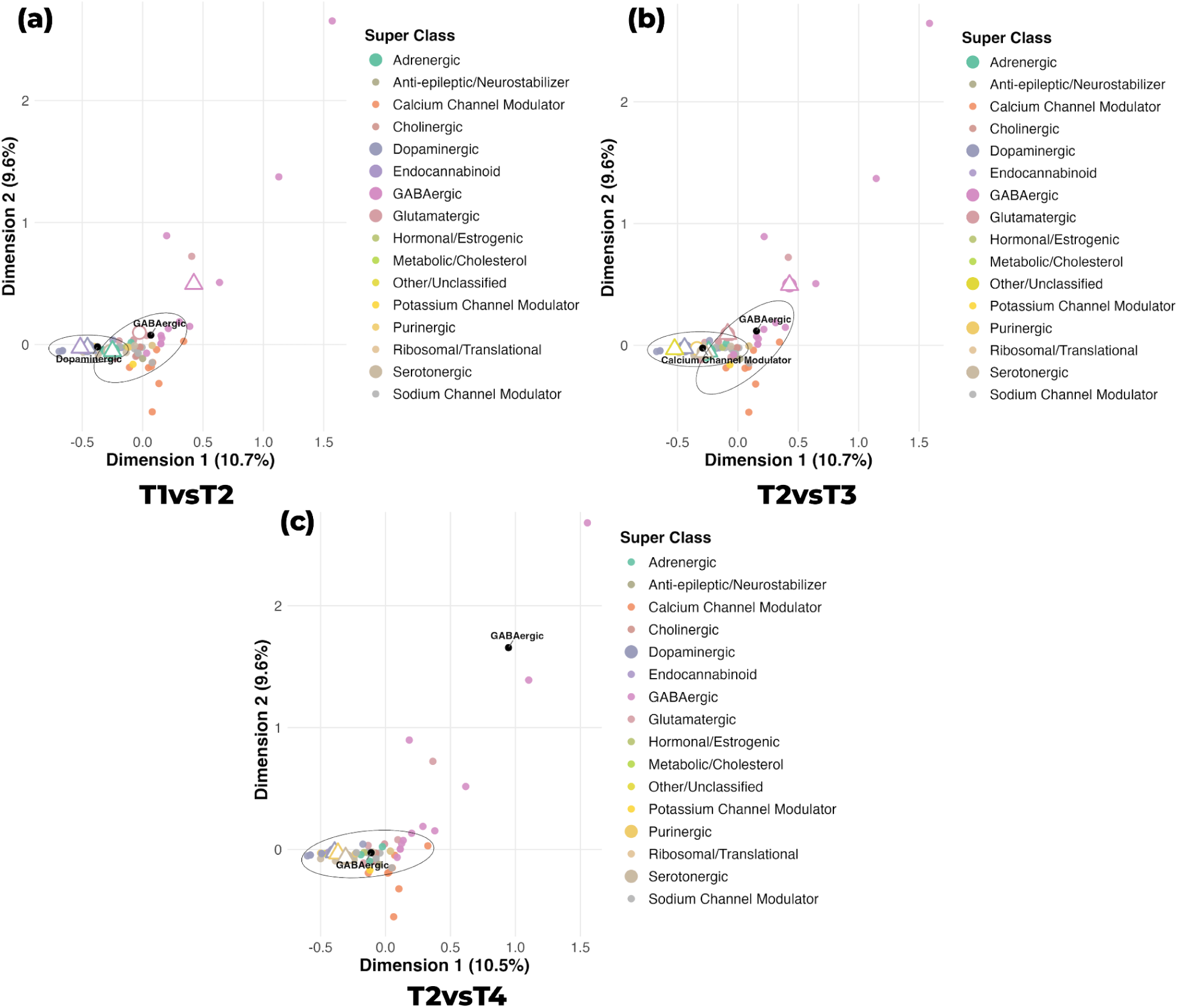
Multiple Correspondence Analysis (MCA) of TMS-drug response profiles, overlaid with CMAP-derived pharmacological signatures reveal temporal trajectories of meditation-related modulation. In all three panels each point represents a drug positioned based on its multivariate TMS excitability profile, presence of drug class in CMAP and colored by super drug class. The colored circles denote TMS-based drug clusters according to their super class. Unfilled white-filled circles represent CMAP-positive matches and unfilled triangles represent CMAP-negative matches, indicating similarity or anti-similarity of gene-expression signatures relative to meditation time-point transitions. The bigger circles in legend represent the presence of these classes. (a) T1vsT2 map shows a compact cluster dominated by GABAergic and dopaminergic drugs, indicating the early-phase changes in TMS metrics and activation of reward axis. Most other classes remain dispersed, suggesting minimal global shifts in neurotransmitter systems at this stage. The CMAP classes that emerged includes Glutametergic, dopaminergic and Seretonergic, Endocannabinoids, Adrenergic with positive and negative connectivity (b) T2vsT3 map shows cluster containing GABAergic and Calcium Channel Modulator drugs, implying coordinated engagement of inhibitory and ion-channel mechanisms during the phase of meditation. The appearance of CMAP-positive and CMAP-negative centroids shows early pharmacogenomic convergence with these classes. (c)The T2vsT4 highlights a distinct cluster dominated by GABAergic drugs. The CMAP shows presence of Seretonergic, Purinergic and Dopaminergic classes. This suggests a later-stage shift involving regulation to maintenance, consistent with longer-term neurophysiological adaptation.

In the post-meditation phase (Figure 2c), GABAergic drugs dominated with overlay to, dopaminergic and serotonergic drug classes from CMAP. This pattern highlights an inhibitory tone alongside modulatory influences in TMS responses. The ellipses in panels Figure 2(a–c) reflect the cluster membership which remains stable across time-points with minor positional shifts suggesting that meditation-induced neurophysiological changes map onto the same core pharmacological axes.

To evaluate the robustness of the MCA-based clustering across three time points, we performed bootstrapping with additional statistical measures. Specifically, the Hopkins statistic, mean silhouette values, and Fisher’s exact test for cluster quality and cohesion (Supplementary table S6). The Hopkins statistic remained consistently high across time points (0.815–0.921), indicating strong non-random cluster structure. Mean silhouette values supported this, showing moderate cohesion for T1vsT2 (0.307) and T2vsT3 (0.274), and higher cohesion for T2vsT4 (0.789), reflecting tighter grouping at this comparison. Fisher’s exact tests to assess associations between clusters and the underlying categorical variables for T1vsT2 (p = 0.00377) and T2vsT3 (p = 0.000324), whereas T2vsT4 showed no significant dependence (p = 0.4699), suggesting less variation in the categorical patterns. Collectively these values indicate that the clustering patterns observed are statistically meaningful. Supplementary Figure 3(a–c) shows the bimodal distribution of MCA coordinates in the latent space and the resulting cluster assignments across bootstrap iterations. In all cases, the mean ARI values were well below 1, indicating that none of the bootstrap iterations reproduced the original clustering perfectly. However, the distributions remain well above 0, showing that the bootstrap solutions do not collapse to unrelated or random clusterings. Instead, they exhibit moderate stability, an expected pattern for categorical, sparse datasets analyzed using MCA.

Augmenting our MCA, we found direct overlap of 5 drugs, belonging to three classes, overlapping with our connectivity data for which their effects on motor system physiology through TMS has been studied (Figure 3). Most striking was Zolpidem which is a GABA- benzodiazepine receptor agonist. This drug is used for treating insomnia. Typically benzodiazepine agonists have been found to increase long-intracortical inhibition (LICI), short-intracortical inhibition (SICI) and short-afferent inhibition (SAI) (refs). Zolpidem has also been found to increase the TEP signature N45 in sensorimotor regions emphasizing on increased early α-synchronization. All these markers represent changes in the GABA-mediated inhibitory system. Moreover since the benzodiazepine agonists showed positive connectivity for T2vsT3, it represents that meditation could have effects that can mimic this drug class and one can expect to see similar changes in the motor suppression post meditation.

**Figure 3:**
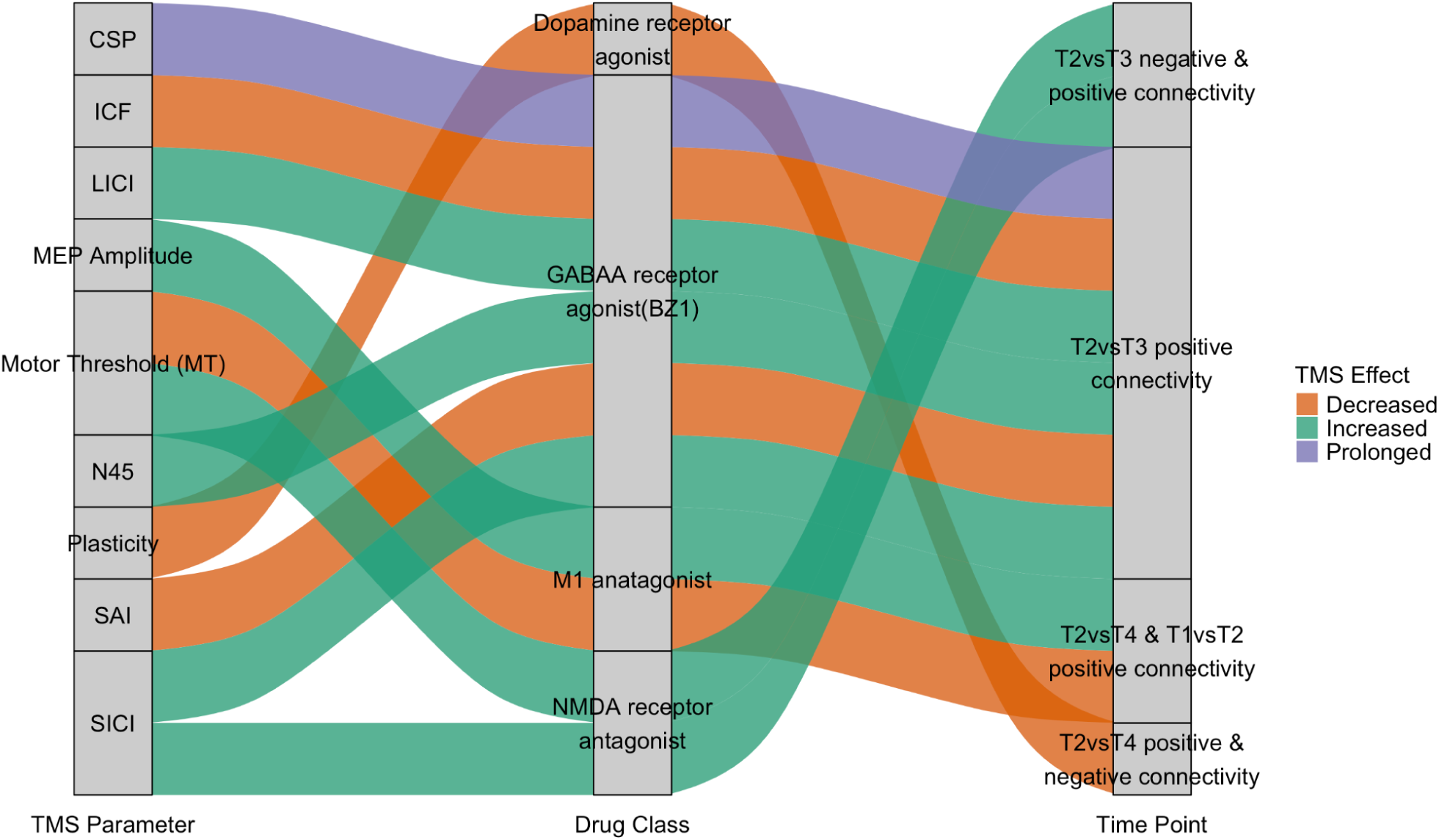
Sankey diagram illustrating the interplay between three axes: pharmacological classes, motor physiology metrics, and temporal connectivity. The diagram shows the connections between three overlapping drugs, grouped by class, and literature-reported TMS responses to these pharmacological agents.These effects were integrated with time-resolved connectivity profiles to examine the progression of neurophysiological modulation across study phases. During the T2 vs T3 meditation interval, benzodiazepine agonists were linked to elevated LICI, SICI, and SAI measures, together with lengthened cortical silent periods, reflecting strengthened inhibitory control. NMDA receptor antagonists display both positive and negative connectivity patterns, influencing SICI and ICF. M1 antagonists show positive connectivity at T2vsT4 (post-meditation) and T1vsT2 (pre-meditation), corresponding to increased MT and SAI and decreased MEP. The color-coded flows represent the direction of TMS responses.

M1 antagonists were another class of drugs (such as **Scopolamine**) for whom their effects on the motor system have been investigated. These drugs increase motor excitability by decreasing the motor thresholds (stimulation intensity for which a muscle twitch is generated is lower) and as a consequence there is an increase in the motor evoked potentials, aka MEPs (refs). Concurrently, the downregulation of these pathways seems to result in a positive connectivity between T2vsT4, suggesting that long-term changes in motor excitability could be a consequence of meditation. Finally, NMDA-receptor antagonists increase SICI and reduce intracortical facilitation (ICF)(16). On the meditation side, the connectivity analysis reveals mixed effects, where both positive and negative connectivity between T2vsT3 was observed, suggesting an overall reduction in the excitatory processes in the brain.

Taken together, the changes in the EI-balance can become a strong marker or mechanism by which meditation can act to lead to behavioral and neurophysiological changes. Our bidirectional approach of examining the similarity at both drug and drug class levels provided a preliminary direction for linking meditation induced molecular changes with its known TMS effects. This framework offers to generate an hypothesis to test the pharmacological-like effects of a non-pharmacological intervention such as meditation within motor system physiology.

## Discussion

The aim of this study was to explore the potential connections between the mechanisms underpinning non-pharmacological meditation techniques and medication targets. By integrating molecular signatures with literature-derived data on drug-induced modulation of cortical excitability, we aim to gain insight into the neurophysiological processes that meditation may influence. A connectivity map study of transcriptomes from 106 participants in an inner engineering meditation retreat revealed strong connections with over 438 medicines used in neuro-psychiatric, cancer, addiction, and hypertension therapies. The clinical study(3,17) had also revealed a number of positive outcomes, including altered levels of cellular anandamide, anti-inflammatory and analgesic reactions, vascular relaxation, and decreased levels of atherosclerosis-associated metabolites that correlate with the drug signatures. Many of the drugs obtained through CMap target GABA, endocannabinoid, dopamine, serotonin, acetylcholine, and epinephrine signaling pathways, which affect behavioral, emotional, and physical states and have been associated to psychosis, anxiety, pain, and inflammation(2,18).The neuroactive ligand receptor interaction pathway was found to be more important during T2 and T3. These pathways correspond to meditation’s claimed benefits on heart rate, stress response, blood pressure, respiratory function improvement, cognitive enhancement, and immune function enhancement(19). Certain pathways, nearly entirely related with T3, are linked to GABAergic synapse, nicotine, and morphine addiction. Our findings are consistent with previous research that links inflammation, immunological response, and neuroactive receptor signaling pathways, providing alternative options for intervention in diseases such as Parkinson’s and multiple sclerosis(20).The absence of medication signatures in the post-meditation phase (T4) emphasizes the significance of consistent practice to preserve therapeutic benefits. The convergence between CMAP-identified drugs and meditation-related signatures motivated us to explore how these pharmacological profiles might relate to changes in brain excitability, via excitation–inhibition (E/I) balance. To establish this connection, we conducted a literature survey of studies employing TMS-EMG and TMS-EEG to characterize drug-induced modulation of cortical excitability. This integrative approach enabled us to contextualize our multi-level findings and to draw mechanistic parallels between meditation-related neurophysiological changes and the effects elicited by established pharmacological agents.

The clustering of drugs grouped by their super drug classes and their alignment with temporal CMap drug classes provided key insights into how meditation may modulate E/I balance. In the preparatory phase, dopaminergic drug classes are prominent, suggesting the reward–motivation axis. During the meditation phase, GABAergic drugs and sodium channel modulators became prominent, an observation consistent with their established role in regulating sodium and chloride fluxes that directly shape cortical inhibition and balance (14,21,22). This observation syncs well with prior findings in (23) where they showed that meditation is associated with enhanced GABAB-dependent inhibitory mechanisms in the cortex, indicating a strengthening of inhibitory tone during practice. Likewise, (24) demonstrated that medications which increase GABAergic activity tend to increase cortical inhibition, whereas those that dampen inhibitory pathways push the system toward greater excitability. In the post-meditation phase, the clusters shifted toward a combination of GABAergic, serotonergic, and dopaminergic classes, reflecting a more integrated neuromodulatory landscape. Conclusively the temporal shifts of meditation converge well onto the pharmacological axes primarily GABAergic, Dopaminergic, and Calcium-channel related mechanisms as independently supported by both TMS drug response literature and CMAP gene-expression similarity.

Altogether, these patterns form coherent pharmacological and neurophysiological patterns that map well onto the temporal changes observed with meditation. This provides insights on how meditation traverses with specific mechanisms of neural excitation and inhibition. Whether we analyse CMap drugs and their associated pathways, or the TMS responses of drugs grouped by their super classes, they consistently converge onto the same fundamental axes of neural modulation. The pathways like nBDNF, calcium signalling, Ras pathways that maintain core processes that impact overall well-being (Figure 1 b) and have been reported repeatedly in other reports (9,23).

These results highlight how meditation induces neurophysiological shifts that mimics modulatory and plasticity-like effects. Recent study using fMRI and blood-plasma analyses report similar outcomes, showing that meditation gives coordinated neural and molecular changes. These changes lead to metabolic reprogramming, and modulation of functional cell signaling pathways (8).

**Figure 3:**
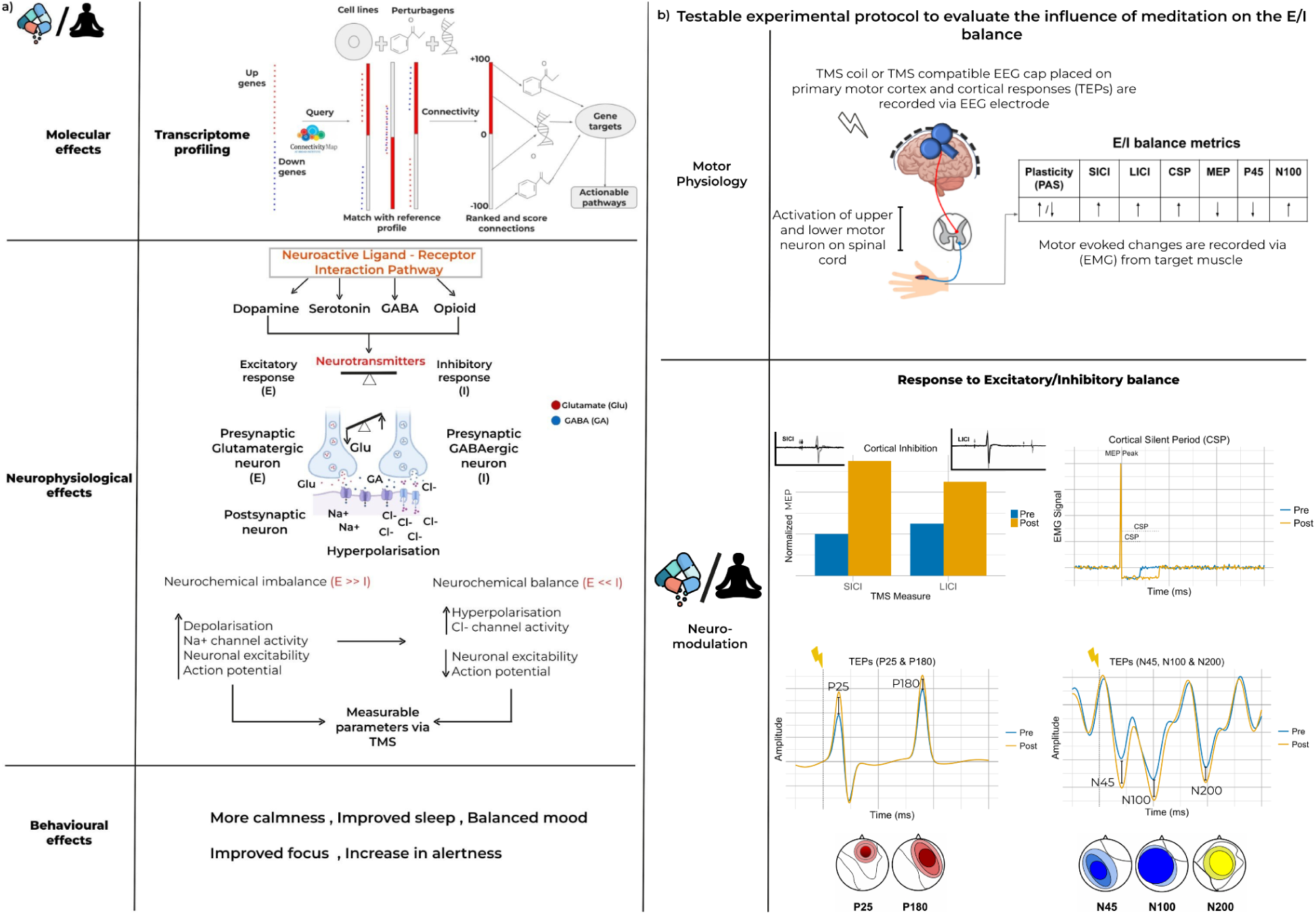
Illustration of the working model for measuring meditation-induced physiological changes across brain regions. **a)** The figure presents a conceptual hypothesis that meditation can induce physiological effects comparable to those of pharmacological agents. At each level—molecular, cellular, and systemic, there is measurable connectivity. At the molecular level, transcriptomic signatures derived from meditation are mapped to the Connectivity Map (CMap; www.clue.io), revealing both positive and negative connectivity with various drug classes. The prominent classes included antipsychotics, depressants, and drugs related to immune and infection. Remarkably, the gene targets of these drugs and meditation appear to converge on common biological pathways, with most overlap via Neuroactive Ligand-Receptor Interaction Pathway. This shared pathway suggests that both drug action and meditation can produce similar neurophysiological effects in specific brain regions. These changes help maintain the brain’s excitation/inhibition (E/I) balance, which can manifest in observable behavioral phenotypes such as increased calmness or enhanced attention. Such changes are also measurable via transcranial magnetic stimulation (TMS). b) highlights the testable protocols which can be used to study the effects of meditation on brain’s physiology using TMS-EEG and TMS-EMG and their associated metrics. A prediction from our model that the post meditation changes should lead to an increase in LICI, SICI and CSP, all of which are GABA-ergic mediated changes. The TMS-evoked potentials will also provide a window into the changes produced post meditation which increase in P25, P180 and decrease in the N25, N100 and N200 potentials. All of these predicted changes are from the drugs which overlap with the connectivity analysis.

Our hypothetical model for the functional effects of meditation builds on these findings. We propose a mulit-level model for the effect of meditation on the molecular, neurophysiological and as a result the behavioral level as seen in Figure 3A. Meditation is proposed to alter first at the transcriptomic level, leading to changes in the neuroactive ligand receptor pathways. This specifically includes the dopaminergic, serotonergic, GABA-ergic and opioid-mediated pathways. These pathways have been implicated in regulating stress and affective balance (25). These changes at the molecular level then translate into neurophysiological effects, specifically affecting the EI-balance of the cortical brain regions. Particularly, the shift in the balance towards excitation (E >> I) is restored via meditation through its potential inhibitory effects (E << I), the nature of which can be measured using parameters measurable via non-invasive stimulation such as TMS.

We propose that these changes can be empirically tested using non-invasive neurostimulation techniques, i.e., TMS combined with EEG/EMG, which provide direct markers of cortical excitability, inhibition (e.g., short/long-interval intracortical inhibition, cortical silent period), and plasticity (Figure 3B). Experimental outcomes pre- and post-meditation showing increase in SICI, LICI and silent period can provide us with the immediate effect of the meditation. On the other hand, long-term plasticity effects can be ascertained using the TMS-evoked potential (TEP) measurements(26). In the end, we predict that this in turn will lead to behavioral changes such as increased calmness, improved sleep and mood. Our integration of the molecular-level, neurophysiological-level and behavioral-level provides a holistic framework to suggest that meditation effects can be quantifiable with changes in these brain-metrics. This leads to the following important implications for meditation and practice and for clinical professionals.

All these results highlight the need for regulating the use of meditation as an interventional therapy. Possibly people who are taking antidepressants and antipsychotics and also meditate may have unwanted side effects that could benefit from recalibration of medication or meditation under clinical supervision. This is consistent with previous reports of adverse consequences, which include physiological, psychological, psychopathological, and spiritual occurrences(27). We also found overlap with drugs used in hypnotherapy, such as scopolamine, and ones that produce euphoria. During meditation, suppressed memories of traumas, as well as accompanying emotional feelings, are occasionally reported(28).

Healthcare professionals who use meditation as an additional therapy should be aware of the possible negative and contraindications(4). Worthwhile to mention, meditation methods were typically taught in a tailored manner by experienced practitioners in ancient times. In contrast, the contemporary era’s internet platforms have facilitated the spread of meditation techniques without cognizance of these practices or intellectual grounding. Our study implies that traditional systems may have evolved to minimize negative consequences and increase benefits. More specifically, meditation (mindfulness-based) can lead to a variety of experiences in people practicing without guidance, and ranging from very positive to very negative including enduring long-lasting functional impairments (29). Individual variability has to be taken into account as there have been reports of vulnerable populations with a history of trauma, bipolarity and psychosis to be more susceptible to adverse reactions of meditation (30). Britton 2019 suggests that meditation should also follow, like all interventional practices, an inverted-U shaped curve where its effects can become harmful when it crosses a limit. The titration protocols built in the traditional systems offered a potential safeguard in terms of the duration, frequency and regulation when these practices would have negative effects. Furthermore, our study adds to this picture by suggesting the potential pharmacological effects of meditation on neurophysiology. Any shifts in these neurophysiological parameters could either be beneficial or detrimental. Modern meditation teachers and psychologists must take into cognizance these factors and develop a clear clinical pathway for the type of meditation, its frequency and duration, the current medication regimen as well as psychological vulnerabilities, and the need for supervision before prescribing meditation for overall well-being. It could thus be highly beneficial, if these practices by patients are also taken into due cognizance in clinical settings.

In conclusion, our study suggests a pharmacological effect of meditation from our convergent mapping of the molecular and neurophysiological changes. Our study highlights the variability in meditation response and emphasizes the need for a personalized approach to achieve optimum benefits. This aligns well with the multi-level testable hypothesis proposed in this study, where changes at molecular and neurophysiological levels jointly shape the behavioural and cognitive outcomes. There are, however, limitations to our findings. Our current insights are limited from a specific meditation approach that needs to be further tested in other practices to make the inferences more generalized. Meditation can take several forms from focussed attention, situational monitoring and recitation. It is not clear if all of these forms will evoke similar changes. The observed heterogeneity in transcriptomic and cortical responses among practitioners further highlights the need for personalized approaches that account for each individual’s baseline physiological state, genetic background, and prior experience. The limited overlap at the individual drug level between the TMS literature and the CMAP database restricted our ability to perform direct drug-to-drug comparisons. CMAP operates at the level of gene-expression similarity and can show both positive and negative connectivity for a given drug class, whereas there are many axes of modulation (for instance E/I balance, cortical plasticity) when it comes to the neurophysiological effects. This study revealed unexpected connections between meditation-induced gene profiles, medication reactions and TMS response, thereby supporting the development of evidence-based clinical applications of meditation.

## Acknowledgements

MM and AJ acknowledge financial support from MOA(Ministry Of AYUSH) for Center of Excellence “AyurTech”(S/MOA/MTM/AA/20210105), IIT Jodhpur.Authors acknowledge Debarka Sengupta, IIIT Delhi, for his critical inputs in analysis.

## Supplementary Information

All study data are included in the article, supplementary tables and supplementary figures. The supplementary data is available at: https://drive.google.com/drive/u/2/folders/1BxiwW9qba9mLfx9RNVmtb12Rp82dA1js

