## Supplementary Figure for "Meditation as a Bioactive Intervention: Molecular and Neurophysiological Mechanisms Revealed by Connectivity Mapping"

### Supplementary Figures

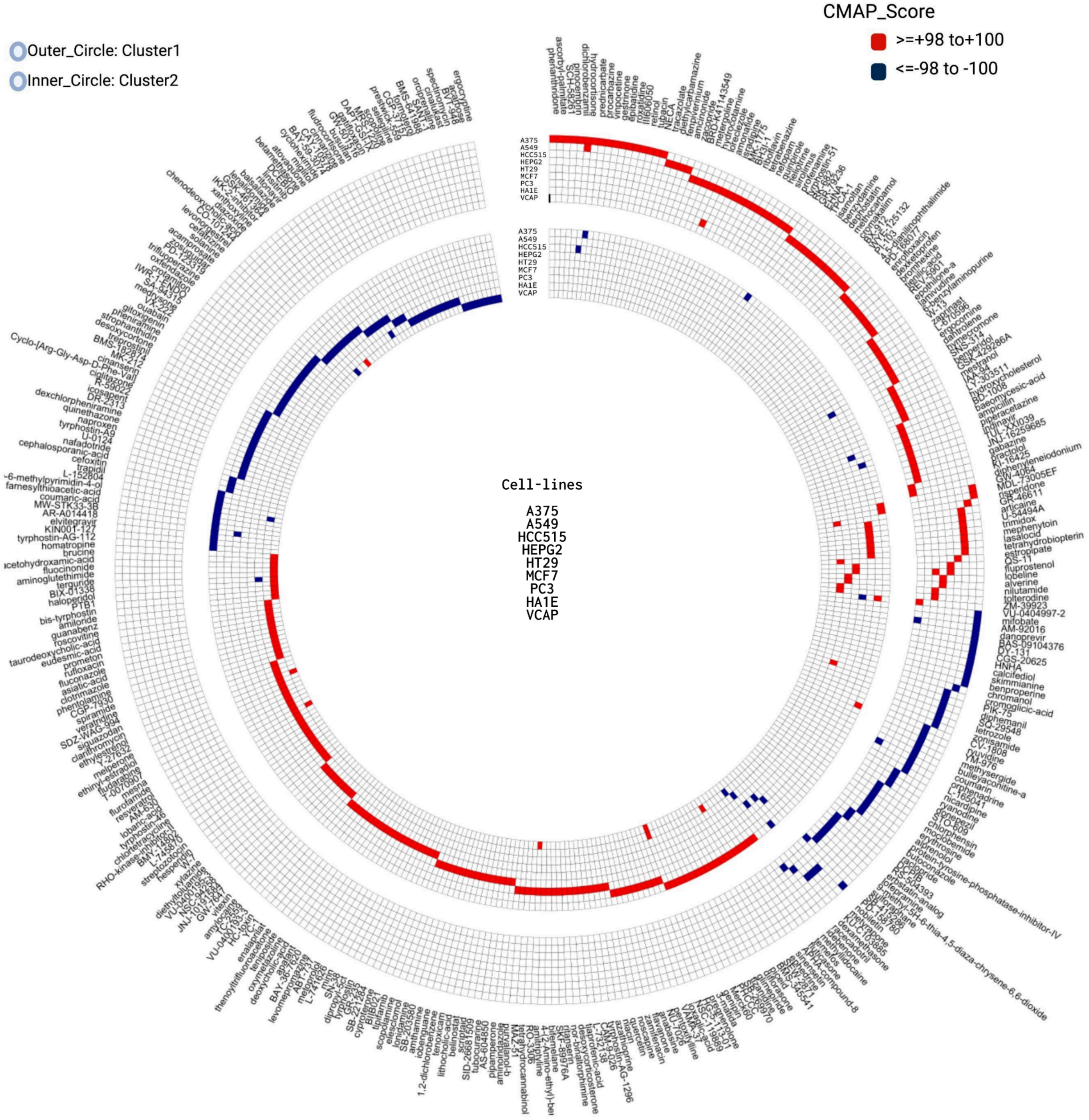


**Figure S1: Circos plot for intersample variability analysis**. The plot shows the difference between the T2 vs T3 time points in terms of drugs from CMAP.This difference highlights that the drugs observed at these time points are merely not the result of cumulative effect of transcriptomic signature overlap with pharmacological agents from CMAP rather it shows the biological difference that exists between the groups.

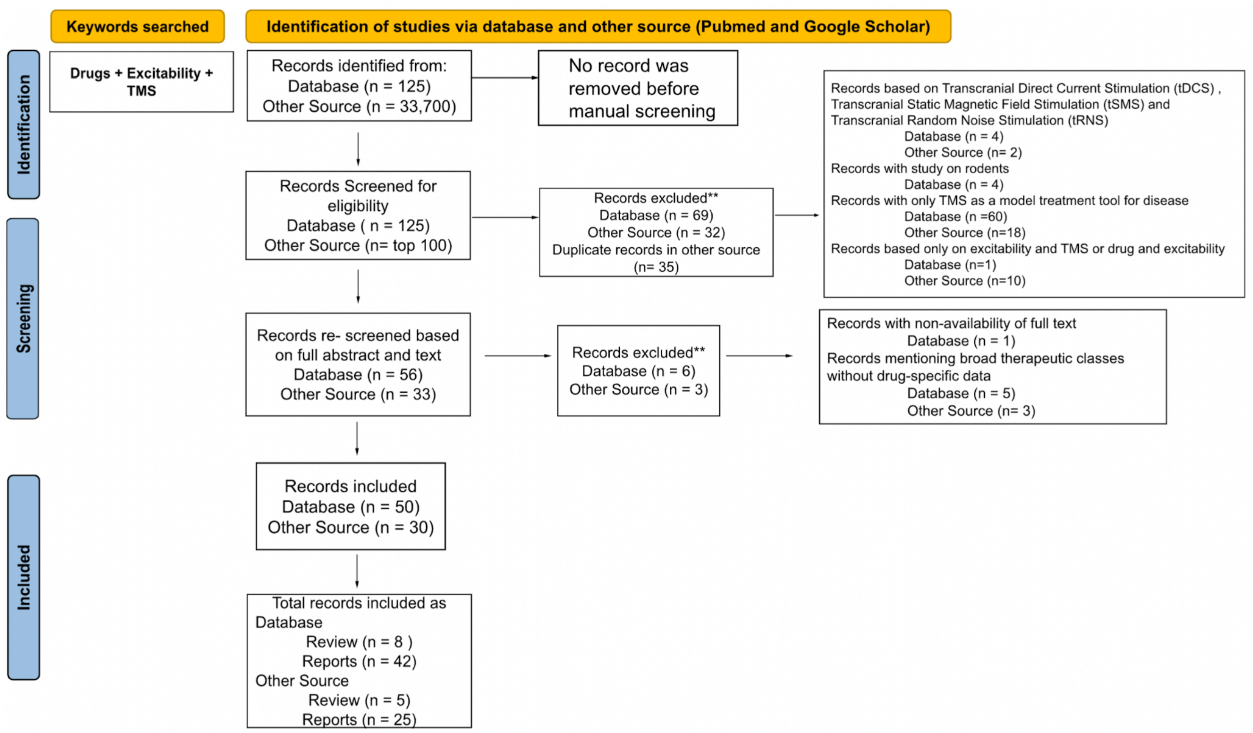


**Figure S2: PRISMA flowchart summarizing the literature search of drugs studied for their cortical excitability via TMS**. The flowchart details the selection process of research articles from Pubmed and Google Scholar using the search keywords "Drugs AND TMS AND Excitability". The duplicate articles were screened between databases. For Google Scholar the top 100 articles were screened. After abstract and full text screening, studies were included if they reported quantitative TMS-EMG/EEG metrics for at least one drug. Based on the inclusion criteria, 50 articles from Pubmed and 30 from Google Scholar were shortlisted. From the final set of studies, a comprehensive table summarizing all drugs, their classes, and corresponding TMS responses was compiled. The references are mentioned at the end of this document.

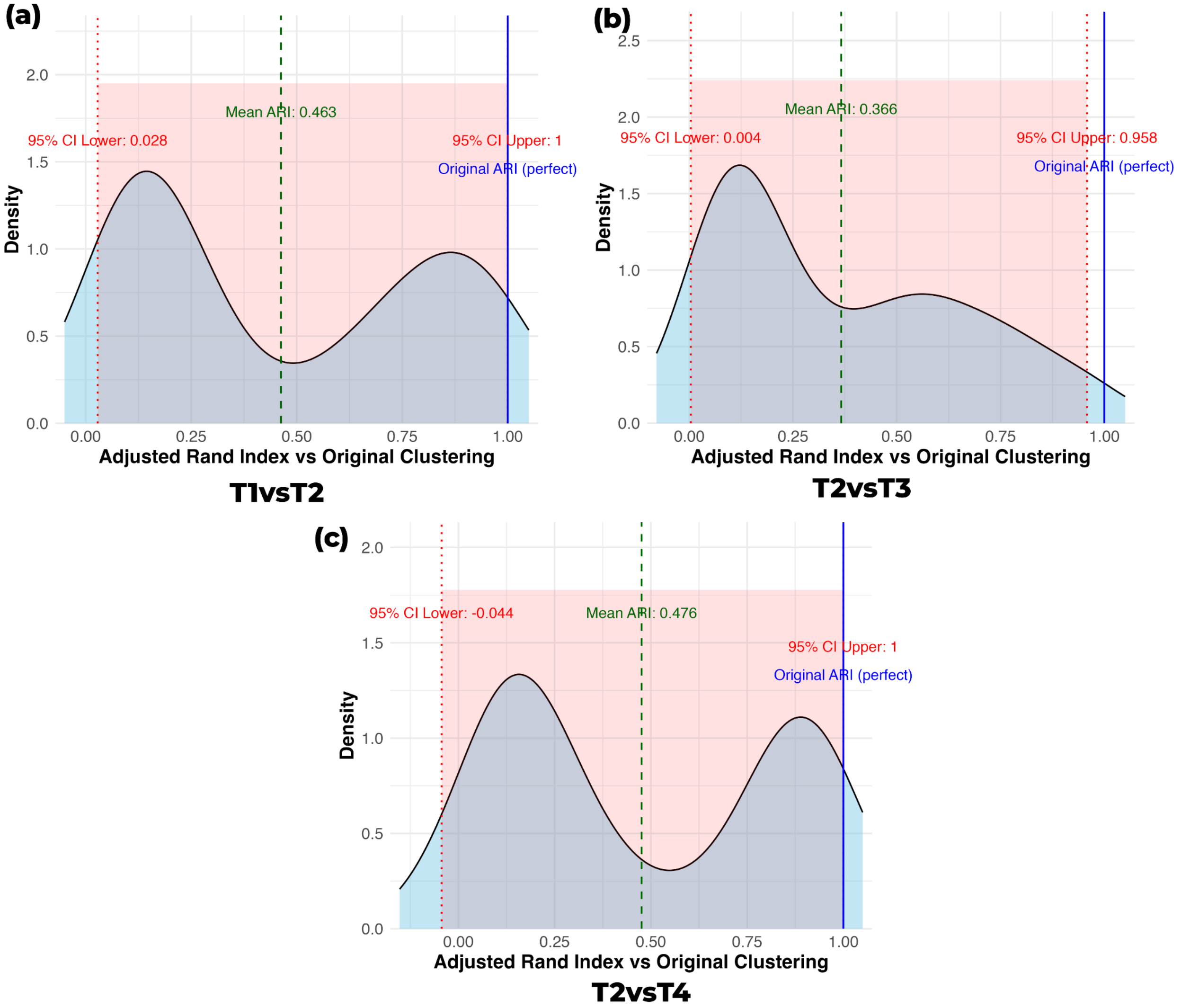


**Figure S3: Bootstrap stability analysis of MCA-derived drug-response clusters across time points.** Panels (a–c) show the distribution of the Adjusted Rand Index (ARI) comparing the original MCA clustering with 300 bootstrap-resampled datasets. The density curves represent how often the bootstrap solutions recover the original cluster structure. The dashed green line marks the mean ARI, dotted red lines indicate the 95% confidence interval (CI), and the solid blue line marks the ARI = 1 reference. The light red shading denotes the extent of the CI. (a) T1 vs T2: The ARI distribution (mean ≈ 0.46; CI ≈ 0.03–1.0) indicates moderate stability.(b) T2 vs T3: The mean ARI (≈ 0.37; CI ≈ 0.00–0.96) suggests moderate stability with greater variability in how consistently the bootstrap samples reproduce the original partitions. (c) T2 vs T4: This comparison shows the highest mean ARI (≈ 0.48; CI ≈ –0.04–1.0). Despite the wide CI, the distribution retains a clear central peak.

66. Uzunova G, Pallanti S, Hollander E. Excitatory/inhibitory imbalance in autism spectrum disorders: implications for interventions and therapeutics. World J Biol Psychiatry.

2016;17(3):174–186.

78. Li X, Tenebäck CC, Nahas Z, Kozel FA, Large C, Cohn J, et al. Interleaved transcranial magnetic stimulation/functional MRI confirms that lamotrigine inhibits cortical excitability

in healthy young men. Neuropsychopharmacology. 2004;29(7):1395–1407.

79. Korchounov A, Ziemann U. Neuromodulatory neurotransmitters influence LTP-like plasticity in human cortex: a pharmaco-TMS study. Neuropsychopharmacology. 2011;36(9):1894–1902.

80. Ziemann U, Tergau F, Bruns D, Baudewig J, Paulus W. Changes in human motor cortex excitability induced by dopaminergic and antidopaminergic drugs. Electroencephalogr Clin Neurophysiol. 1997;105(6):430–437.
